## Supplemental Methods for "From Synapses to Dynamics: Obtaining Function from Structure in a Connectome Constrained Model of the Head Direction Circuit"

### A Technical Appendices and Supplementary Material

#### A.1 Weight Symmetrization

To enforce bilateral symmetry in the neural network architecture while preserving functional connectivity patterns, we implemented a weight matrix symmetrization procedure. Let $\mathbf{W} \in \mathbb{R}^{N \times N}$ be the initial weight matrix, where $N$ is the number of neurons. For each neuron $i$, we define its lateralization $l_i \in \{L, R\}$ and its lateralized label $\tilde{l}_i$ (removing lateralization information).

The symmetrization process operates on connection groups defined by the tuple $(s, t, l_s, l_t)$, where $s$ and $t$ are the lateralized labels of the source and target neurons, and $l_s$ and $l_t$ are their lateralizations. For each unique connection group, we compute the symmetrized weight:

$$w_{sym}(s, t, l_s, l_t) = \frac{1}{2n_{st}} \sum_{i,j} w_{ij} \mathbb{I}[(s, t, l_s, l_t) = (\tilde{l}_i, \tilde{l}_j, l_i, l_j)]$$

where $n_{st}$ is the number of connections in the group and $\mathbb{I}$ is the indicator function. The final symmetrized weight matrix $\mathbf{W}_{sym}$ is constructed as:

$$W_{sym,ij} = \begin{cases} w_{sym}(\tilde{l}_i, \tilde{l}_j, l_i, l_j) & \text{if } \exists w_{sym}(\tilde{l}_i, \tilde{l}_j, l_i, l_j) \\ W_{ij} & \text{otherwise} \\ 0 & \text{if } i = j \end{cases}$$

This procedure ensures that left-right symmetric connections have identical weights ($W_{sym,ij} = W_{sym,ji}$ for lateralized pairs), while preserving the network's functional connectivity patterns and maintaining the overall network structure and strength. The result of this symmetrization is shown in Figure A1.

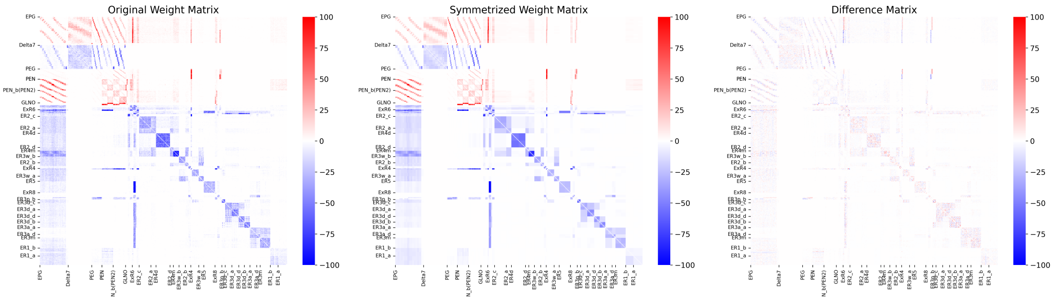


Figure A1: Weight matrix symmetrization.

#### A.2 Single parameter and E/I controls

In addition to the cell-type parameterization of the model, we also explored using a single parameter to describe the synapse gains and a parameterizing the excitatory and inhibitory neurons separately. These reduced parameterizations fail to produce a ring attractor following training (Figure A2).

#### A.3 Kernel PCA and loss landscape visualization

In order to visualize the optimization landscape, we ran training runs with differently seeded initial parameters. We tracked the parameter values throughout training. We then used kernel PCA (Schölkopf et al., 1998) with a radial basis function kernel to fit the training trajectories and used a Kernel Ridge regression to learn the mapping from the basis to the original parameter space which we use to sample parameter values on in order to construct a two dimensional loss landscape.

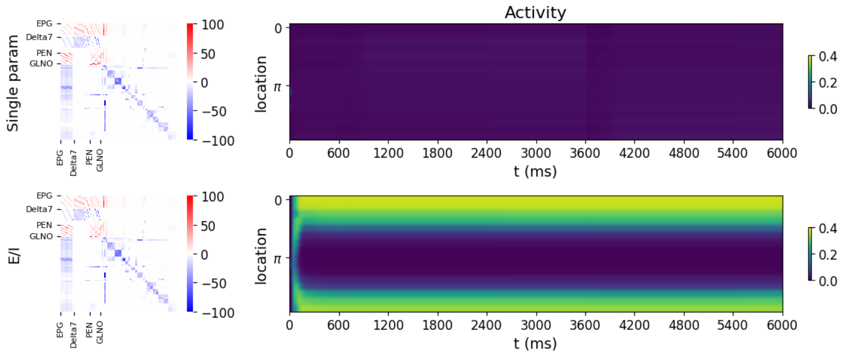


Figure A2: Controls for optimizing the network with a single global gain or gain parameters for the excitatory and inhibitory neurons.

##### A.4 Hyperparameter sweeps and model selection

For each ablation experiment (including the no-ablation condition), we trained models across a set of 90 hyperparameters by using a grid search over the initial bias values, leak, and global synapse strength. We additionally randomly initialized the block weights using a normal distribution centered at 1 with variance of 2 percent of the parameter value and swept over five seeds for each hyperparameter setting. In Figure 5 we chose the best 5 models from each condition to evaluate. We define best as the models with the smallest minimum speed loss (i.e., the models with the greatest range of velocity integration), after filtering for a R2 of $> 0.7$ and a temporal consistency loss of $<$ 0.013.

##### A.5 Delta7 ablations with full ring parameterization

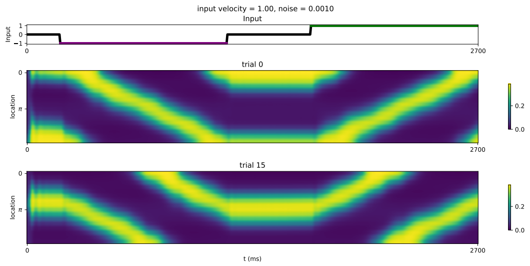


Figure A3: Activity bumps in Delta7 ablation, full ring parameterization.

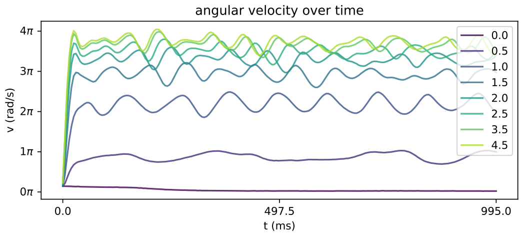


Figure A4: Internal vs. input velocity in Delta7 ablation, full ring parameterization.

#### A.6 Sample weights from ablation experiments

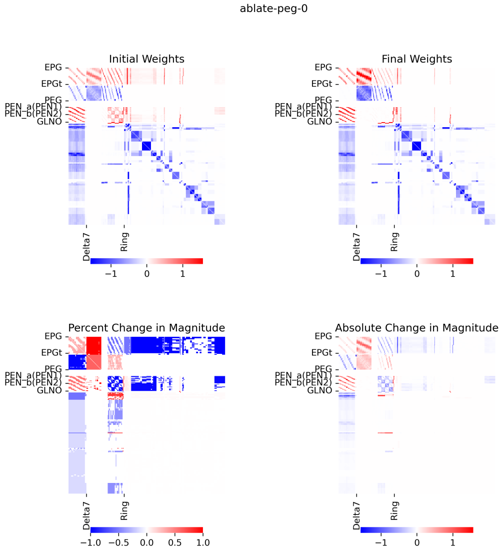


(a) P-EG ablation

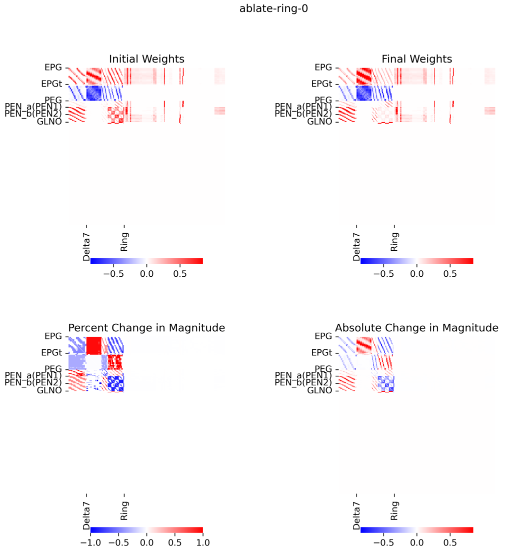


(b) Ring ablation

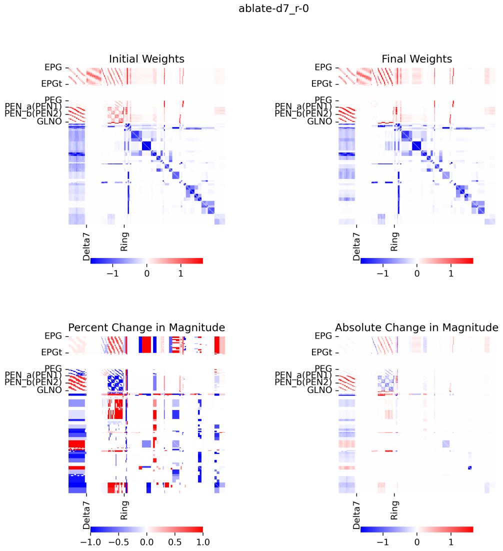


(c) Delta7, full ring parameterization

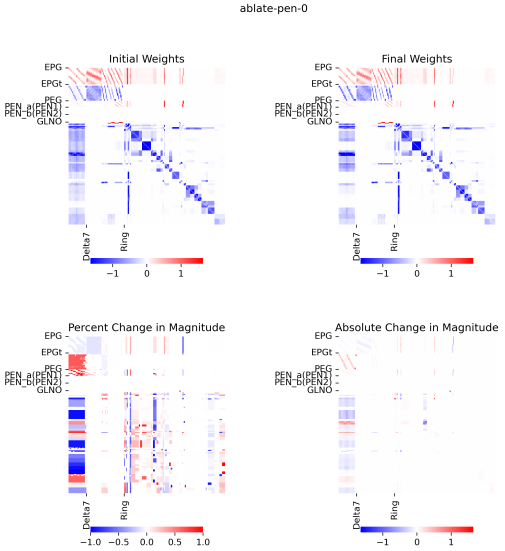


(d) P-EN ablation

Figure A5: Ablation effects on trained synaptic weights: P-EG, Ring, Delta7, and P-EN.

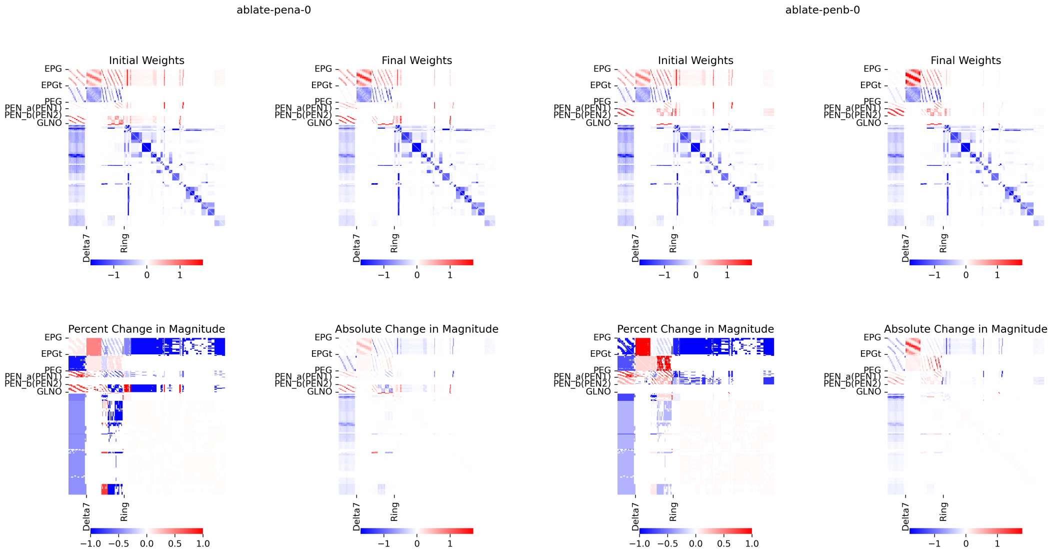


(a) P-ENa ablation

(b) P-ENb ablation

Figure A6: Ablation effects on trained synaptic weights: P-ENa and P-ENb.

#### A.7 Additional Supplementary Methods

**Velocity input** Velocity input is driven by differential activity between the left and right GLNO neurons. In our experiments, we drove the velocity using either the left or right GLNO neurons at a given time. It is sufficient to drive differential activity between the left and right GLNO neurons to drive the velocity input which can be shown in A7.

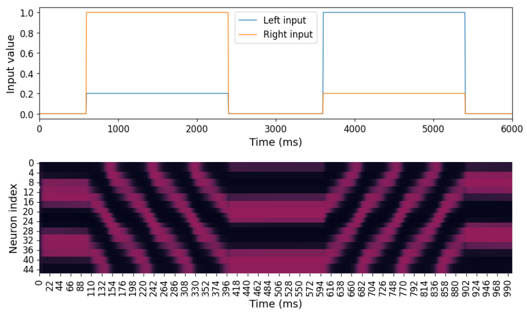


Figure A7: Activity trace when reversing sign of velocity input

#### A.8 Supplementary Figures

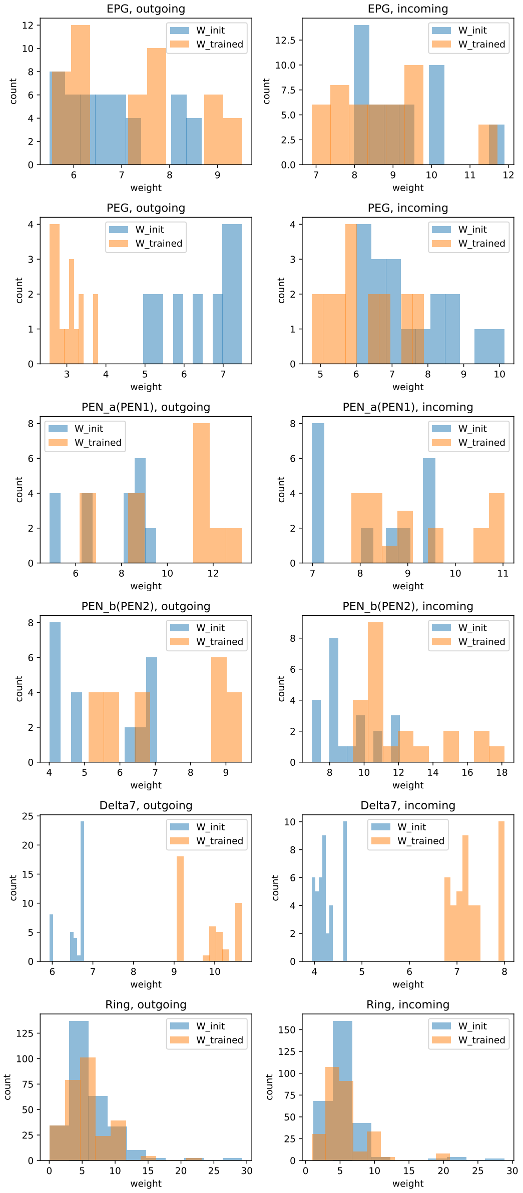


Figure A8: **Synapse statistics.** Distribution of incoming and outgoing synapse weights before and after training of a full connectome-initialized network.
